# Molecular Insights on exquisitely Selective SrtA inhibitors towards active site loop forming open/close lid conformations in SrtA from *Bacillus anthracis*

**DOI:** 10.1101/710251

**Authors:** Chandrabose Selvaraj, Gurudeeban Selvaraj, Satyavani Kaliamurthi, Dong-Qing Wei, Sanjeev Kumar Singh

## Abstract

The present study clearly explains the dependency of inhibitory activities in SrtA inhibitors is closely related to protein conformational changes of SrtA from *Bacillus anthracis B. anthracis*Sortase A (SrtA) protein anchors proteins by recognizing a cell wall sorting signal containing the amino acid sequence LPXTG In order to analyze conformational changes and the role of SrtA enzyme, especially the loop motions which situated proximal to the active site molecular dynamic simulation was carried out for 100ns. Particular loop is examined for its various conformations from the MD trajectories and the open/close lid conformations are considered for the enzyme activity validations. Experimentally verified SrtA inhibitors activity was analyzed through 3D-QSAR and Molecular docking approaches. Results indicate that, biological activity of SrtA inhibitors is closely related to the closed lid conformation of SrtA from *Bacillus anthracis*. This work may lead to a better understanding of the mechanism of action and aid to design a novel and more potent SrtA inhibitors.

## Introduction

*Bacillus anthracis*, a gram-positive, spore-forming bio-threat bacterium that is the etiological pathogen of the zoonotic disease anthrax. It constantly exposed to different environmental conditions in its host and strongly influencing its physiology. It is a life-threatening infectious disease, widespread in most parts of the globe, and cases of anthrax have been reported from almost every country. Despite the emphasis on its role as an agent of bioterrorism or biological warfare, anthrax is continuing to be, an important global disease of wildlife and livestock^**1**^.Currently, novel approaches are required for the treatment of bacterial infections, due to continuous changes in the patterns of infectious disease and the emergence antibacterial resistant to present antibiotics. Currently, researchers are involved in finding drug targets of great interest in terms of understanding and inhibiting the infection process and these targets may have potential as targets for therapy^**2**^.

Surface proteins of this bacterium play an important role in host-pathogen interactions and this mechanism have attracted theresearchersin the hunt for the drug target to inhibit the pathogenic microbes. Cell wall anchoring occurs through a transpeptidation mechanism, which requires surface proteins with C-terminal sorting signals^**3**^. In sortase family of proteins, class A enzymes have attracted significant interest as a universal drug target of all Gram positive pathogens and the clinically important Gram positive pathogens use the Sortase A (SrtA) gene to display virulence factors. SrtA enzymes are playing a vital role incleaving the sorting signals of secreted proteins to form isopeptide (amide) bonds between the secreted proteins and peptidoglycan or polypeptides^**4**^. SrtAis involved in the covalent anchoring of surface protein to the cell wall peptidoglycan, or link proteins together to construct pili which promote the bacterial adhesion to host cell and tissue to acquire the essential nutrients. These enzymes are actively involved as the principal architects of the bacterial surface that makes SrtA as an attractive drug target^**5**^.

Recent research has came up with evidence, that virulence of gram positive pathogenic microbes lies on SrtA enzymes, and removal of SrtA gene shows lack of pathogenicity and cell wall formations^**6**^. This evidence has enlightened and attracted the researchers to develop the promising lead considering SrtA as a drug target. SrtA, is main focus of this study, anchors seven proteins to the cell wall by joining the threonine of the LPXTG sorting signal at the C-terminus of the protein to the amine group of meso-diaminopimelic acid (DAP) within lipid II^**7**^. In this mechanism, the key active site loop (β7/β8 loop) undergoes a disordered-to-ordered transition upon binding the sorting signal, potentially facilitating recognition of lipid II.Hydrophobic residues Val168 and Lue169 located in β6-β7 loop which involved hydrophobic interaction with N-terminal LPETG peptide and the catalytic residues Pro168, Asp169, Lys170 and Trp171 are present in the loop region between β7/β8^**8**^. Our studies focuses on structural changes in SrtA active sites loops (β7/β8) and analyze the activity of known SrtA inhibitors against this various loop conformation^**9**^. Suree*et al*.^**10**^ had performed structure–activity relationship (SAR) analysis, which leads to the identification of several rhodanine, pyridazinone and pyrazolethione compounds with more potential activity against the *B. anthracis*thanany other previously mentioned natural or synthetic inhibitors. In this study, we have evaluated the activity of the reported compounds through 3D QSAR model and the active conformations are cross validated with open and closed conformation of β7/β8 loop in SrtA.This study on analyzing the available SrtA inhibitor’s against the β7/β8 loop conformation of SrtA willbe much more helpful in discovering of novel lead molecules for further development.

## Materials and Methods

### Ligand preparation and Pharmacophore Generation

Experimentally active 28 compounds of rhodanine, pyridazinone and pyrazolethione derivatives reported by Suree et al.^**10**^ were taken for the study and the ligand conformations wereprepared by using LigPrep/Confgen with OPLS AA. Each compound structure is represented by a set of points in 3D space, which coincide with various chemical features that may facilitate non-covalent bonding between the compound and its target receptor. PHASE (PHASE V 4.2) provides a built-in set of six pharmacophore features, hydrogen bond acceptor (A), hydrogen bond donor (D), hydrophobic group (H), negatively ionizable (N), positively ionizable (P) and aromatic ring (R). The rules that apply to map the positions of pharmacophoric sites are known as feature definitions, and are represented internally by a set of SMARTS patterns^**11**^. Each pharmacophore feature is defined by a set of chemical structure patterns. All user-defined patterns are specified as SMARTS queries and assigned one of three possible geometries, which define the physical characteristics of the site like point, vector and group. Active and inactive thresholds of pIC_50_ 5.9 and 4.3 were applied to the dataset to yield eight actives and five inactives, which are used for pharmacophore modeling and subsequent scoring.

### Building the 3D-QSAR model

All the generated common pharmacophore hypotheses were used to generate atom-based 3D-QSAR models by correlating the actual and predicted activity for the set of 21 training molecules using PLS analysis. The PLS regression was carried out using PHASE with a maximum of N/3 PLS factors (N=number of ligands in the training set, and a grid spacing of 1.0 Å) and all the generated models were validated by predicting activity of 7 test set molecules^**12, 13**^.

### External validation

In order to understand the predictive abilities of established model was evaluated to confirm the robustness by external validations^**14**^. The calculated parameter is r^2^, K values and 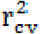. The r^2^ value calculated by the following formula,

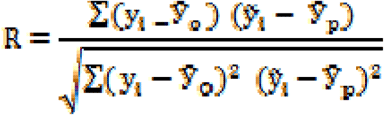

Where, *y*_*i*_ and 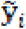 are the observed and predicted activity,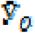 and 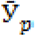 are the average values of the observed and predicted pIC50 values of the test set molecules. For the ideal QSAR model, the r^2^ value should be close to 1. And the regression of y against 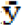 through the origin: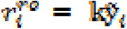 should be characterized by k close to 1. Slope k is calculated as follow:

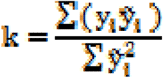

The cross validated co-efficient,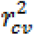, was calculated using the following equation:

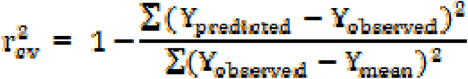

Where, Y_predicted_, Y_observed_ and Y_mean_ the predicted, observed, and mean values of the target property (pIC50), respectively^**15**^.

### Applicability Domain

The applicability domain (AD) prediction was performed by using the SIMCA-P 13.0 demo version (Umetrics. 2002). The AD is the physico-chemical information based on a training set of the model and it’s also applicable to make predictions for new hit compounds. By this approach, one can directly analyze the properties of distance matrix, multivariate descriptor to investigate the applicability domain of the training set. These approaches define the AD by calculating distances of a query compound from a defined point within the descriptor space of the training data^**16**^. This measured distance between defined point and the dataset, then compares with a pre-defined threshold. These calculations are done by feature selection and principal component analysis (PCA). The reliable prediction of the source compounds is likely to be similar to the data set and the activity prediction errors were avoided by using activity prediction applicability domain concept^**17**^.

### Molecular Dynamics Simulation

Here, MD simulation was performed for SrtA structure and also for ligand bound SrtA structures. For the Apo structure of SrtA, the NMR structure of *B. anthracis*SrtA (PDB ID: 2KW8 (http://www.rcsb.org) was subjected to MD simulation for 100ns of timescale, in order to understand the nature of intra-molecular conformational changes of protein structureMD simulation is performed with GROMACS program package (http://www.gromacs.org) adopting the GROMOS96 force field parameters^**18**^. The structure was solvated using the SPC water model, and the solvated structure was energy minimized using steepest descent method, and it’s terminated when maximum force is found in smaller than 100 kJ mol^-1^ nm^-1^. For equilibration and production runs, the temperature was controlled through Nosé–Hoover thermostat dynamics with a damping coefficient of 2 ps^-1^. The long range electrostatic interactions were computed by particle-mesh Ewald method and van der Waals (VDW) cut-off were set to 9 Å. The structure was energy minimized to eliminate bad atomic contacts and subsequently solvated with water. The total simulations were carried out in the NPT ensemble at constant temperature (300 K) and pressure (1 bar), with a time step of 2fs. NVT were performed for 1ns, and the minimized structure was equilibrated with a timescale of 100ns^**19, 20**^. For the SrtA bound ligand complex, receptor and ligands were cleaned using GROMOS96 force field and then the topology files were generated for the receptor and ligands separately using PRODRG tool^**21**^. The simulation system was created manually by importing the ligand topology into the system pursued along with a dodecahedron box with a margin of 1 nm and the system was filled with water using the SPC explicit solvation model. The system was applied with energy minimization and the atomic velocities were adjusted according to Maxwell Boltzmann distribution at 300K with a periodic scaling of 0.1ps. A pre-simulation run of 20 ps was applied to relax the system and to remove the geometric restrains which eventually appeared at the initialization of the run^**22**^. All the simulations were carried out at constant pressure and temperature (NPT) ensemble. The Berendsen coupling was employed to maintain a constant temperature of 300 K and constant semi-isotropic pressure of 1 bar with coupling time of 2.0 fs and the coordinates were saved. The simulation timescale for ligand bound form is 20ns and the RMSD analysis has been performed for understanding the stability of ligands.

### Molecular Docking Simulation

For the interactions, Quantum Polarized Ligand Docking (QPLD), molecular docking, was performedwith open and closed active site. Two different conformation of open and closed active site from the MD simulation are subjected to grid generation. To soften the potential for non-polar part of the receptor, Van-der Waal radii of receptor atoms were scaled by 1.00Å with a partial charge cutoff 0.25^**23,24**^. Ligands are set to be rigid conformations, as of all the conformations from 3D QSAR bioactive conformations were used for these docking studies. The previously reported active site which located around the loop region considered as binding pocket^**25**^.For the specificity of ligand partial charges, QM/MM based QPLD docking was also applied for both the protein conformations with all 28 compounds. For QM calculations, the accuracy level is held in reserve as accurate and background of this calculation was done by using 6-31G*/B3LYP density functional theory^**26,27**^. The charges were calculated for a free ligand by Jaguar, with the presence of the hybrid water model. Here, fixed charges of ligands obtained from force field parameter are replaced by QM/MM calculations in protein atmosphere and only ligand treated as the quantum region. However, the protein ligand complex charge correctness occurs through the QPLD docking and it will enhance the accuracy of results^**28**^.

### Conformation Based Activity Prediction

The Liaison program is an application for estimating the binding affinities between ligands and receptors, using a linear interaction approximation (LIA) model^**29**^. The LIA model is an empirical method fitted to a set of known binding free energies. Liaison runs molecular mechanics (MM) simulations of the ligand-receptor complex, and for the free ligand and receptor using the surface generalized Born (SGB) continuum solvation model^**30**^. The simulation data and empirical binding affinities were analyzed to generate the Liaison parameters, which are subsequently used to predict binding energies for other ligands with the same receptor. The empirical function used by Liaison for the prediction of binding affinities is as follows.

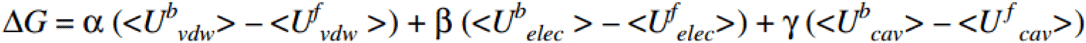

Here, U_vdw_, U_elec_, and U_cav_ are the van der Waals, electrostaticand cavity energy terms in the Surface Generalized Born (SGB) continuum solvent model. Default options were specified including a minimization sampling method using a truncated Newton algorithm. Ensemble averages of van der Waals, electrostatic, and cavity (solvent exposed ligand surface area) energies were computed for the docking conformation with open and closed lid conformations using an implicit solvation model. The computed energies for each inhibitor complex and corresponding pIC_50_ (Experimental Value) were then imported into Strike, where the partial least squares (PLS) and multiple linear regression (MLR) methods were applied, to construct a linear equation representing binding affinity^**31**^.

## Results and Discussion

### Determination of Pharmacophore and 3D-QSAR models

In this study, the experimental active 28 compounds of rhodanine, pyridazinone and pyrazolethione derivatives were used to predict theoretical activity of the test compounds, which are shown in Table 1 and their 2D structure of the scaffoldligand rearrangements are shown in Supplementary figure 1. The biological activity data were reported in the form of IC_50_ values and those values were converted into pIC_50_ using the formula (pIC_50_= -log IC_50_) and the data set was divided randomly into 21 training set and 7 test set compounds in order to maintain the 3:1 ratio^**25**^.We ensured that the structurally different compounds were distributed uniformly with a wide range of pIC_50_ value in both training and test set. Initial step is tend towards generating the pharmacophore model using a tree-based partition algorithm and two best common pharmacophore hypotheses AAAHR, and AAADR were selected for 3D-QSAR model building by applying the scoring function for five featured Pharmacophore hypotheses with default values. Training set compounds were aligned on these common pharmacophore hypotheses and are analyzed in PHASE with three PLS factors and the predictivity of each hypothesis was re-evaluated by the test set compounds. A summary of the statistical data for the two common pharmacophore hypotheses AAAHR and AAADR was listed in Table 2.

**Table 1.** Structures and actual versus predicted pIC50 of compounds (* Test Set Compounds)

| Compounds | X | R1 | R2 | R3 | R4 | Actual | Predicted | Residual |
| --- | --- | --- | --- | --- | --- | --- | --- | --- |
|  |  |  |  |  |  | IC<sub>50</sub> | IC<sub>50</sub> |  |
| Rhodanine |  |  |  |  |  |  |  |  |
| 1-1 |  | -Ph | 4-NO<sub>2</sub> | 3-Br, 2-OH,<br>5-NO<sub>2</sub> |  | 4.69 | 4.52 | 0.179 |
| 1-2 |  | -Me |  | 3-Br, 2-OH,<br>5-NO<sub>2</sub> |  | 4.88 | 5 | -0.11 |
| 1-3 |  | -Pr |  |  | -H | 4.27 | 4.29 | -0.01 |
| 1-4 |  | -Et |  |  | 2-NO<sub>2</sub> | 4.13 | 4.31 | -0.17 |
| 1-5 |  | -Allyl |  |  | -H | 4.56 | 4.52 | 0.04 |
| Pyridazinone |  |  |  |  |  |  |  |  |
| 2-1 * |  | -SH | -OEt | -Ph | -H | 5.85 | 5.88 | -0.02 |
| 2-5 |  | -OMe | -SH | -Ph | -H | 5.74 | 5.5 | 0.24 |
| 2-6 |  | -OH | OCH<sub>2</sub>Ph | -Ph | -H | 4.30 | 3.9 | 0.40 |
| 2-7 |  | -OH | -OMe | -Ph | -H | 4.30 | 4.24 | 0.06 |
| 2-8 |  | -OH | -SEt | -Ph | -H | 4.30 | 4.59 | -0.28 |
| 2-9* |  | -SH | -SEt | -Ph | -H | 6.52 | 6.11 | 0.41 |
| 2-10 |  | -SH | -OEt | -Ph | -H | 5.49 | 5.87 | -0.37 |
| 2-11 |  | -SH | -OEt | -Ph | 4-NO<sub>2</sub> | 5.17 | 5.41 | -0.23 |
| 2-13 |  | -SH | -OEt | -Ph | 3-F | 5.74 | 5.7 | 0.04 |
| 2-14* |  | -SH | -OEt | -Ph | 3-Me | 5.77 | 5.68 | 0.09 |
| 2-15 |  | -SH | -OEt | -Ph | 3,5-Cl<sub>2</sub> | 4.85 | 5.5 | -0.64 |
| 2-16 |  | -SH | -OEt | Cyclohexyl |  | 5.85 | 5.96 | -0.10 |
| 2-17* |  | -SH | -OEt | -Ph | -H | 5.92 | 5.94 | -0.01 |
| 2-18 |  | -OEt | -SH | -Ph | -H | 5.92 | 5.66 | 0.26 |
| 2-19* |  | -OEt | -SH | -Ph | 3-F | 6.04 | 6.16 | -0.11 |
| 2-20 |  | -OEt | -SH | -Ph | 3-Me | 6.39 | 6.25 | 0.15 |
| 2-21 |  | -OEt | -SH | -Ph | 3,5-Cl<sub>2</sub> | 5.28 | 5.48 | -0.19 |
| 2-35* |  | -OEt | -Cl | -Ph | -H | 6.52 | 5.26 | 1.26 |
| 2-36 |  | -OEt | -Cl | -Ph | 4-NO<sub>2</sub> | 3.60 | 4.04 | -0.43 |
| 2-47 |  | -Cl | -Cl | -Ph | 3,5-Cl<sub>2</sub> | 4.85 | 4.71 | 0.14 |
| **Pyrazolethione** |  |  |  |  |  |  |  |  |
| 3-9* | S | 4-N=N-Ph | H |  |  | 5.85 | 5.85 | 0.004 |
| 3-17* | S |  |  |  |  | 6.52 | 6.72 | -0.19 |
| 3-12 | S | 2,4,6-Br<sub>3</sub> | H |  |  | 5.85 | 5.9 | -0.05 |

**Table 2.** Quantitative Structure Activity Relationship (QSAR) results for the two best Common Pharmacophore Hypotheses

|  | AAAHR | AAADH |
| --- | --- | --- |
| SD | 0.34 | 0.41 |
| R<sup>2</sup> | 0.87 | 0.82 |
| F | 40.3 | 25.4 |
| P | 6.079e-08 | 2.555e-06 |
| RMSE | 0.2637 | 0.4111 |
| Q<sup>2</sup> | 0.74 | 0.6806 |
| Pearson R | 0.88 | 0.96 |

**Table 3.** scoring parameters of active compounds in QSAR towards closed lid structure of SrtA

|  | **pIC<sub>50</sub>** | **IFD score** | **Glide score** | **Glide Energy** | **Binding Energy** | **Hydrophilic interactions** |
| --- | --- | --- | --- | --- | --- | --- |
| Pyridazinone-2-19 | 6.046 | 489.21 | -6.591 | -42.62 | -38.61 | Met56, Val110 and Trp171 |
| Pyridazinone-2-9 | 6.523 | 480.67 | -6.486 | -43.19 | -40.51 | Met56, Val110 and Trp171 |
| Pyrazolethione-3-12 | 6.523 | 477.99 | -6.209 | -39.57 | -38.67 | Met56, Val110 and Trp171 |
| Pyridazinone-2-5 | 5.745 | 481.21 | -5.916 | -36.24 | -39.24 | Met56, Val110 and Trp171 |
| Pyridazinone-2-20 | 6.398 | 467.26 | -5.858 | -40.29 | -39.98 | Met56 and Val110 |
| Pyridazinone-2-16 | 5.854 | 459.38 | -5.614 | -36.21 | -38.21 | Met56, Val110 and Trp171 |
| Pyridazinone-2-35 | 6.523 | 461.50 | -5.512 | -38.29 | -38.20 | Met56, Val110 and Trp171 |
| Pyridazinone-2-17 | 5.854 | 446.55 | -5.367 | -39.18 | -36.19 | Met56, Ala58, Val110 and Trp171 |

**Figure 1:**
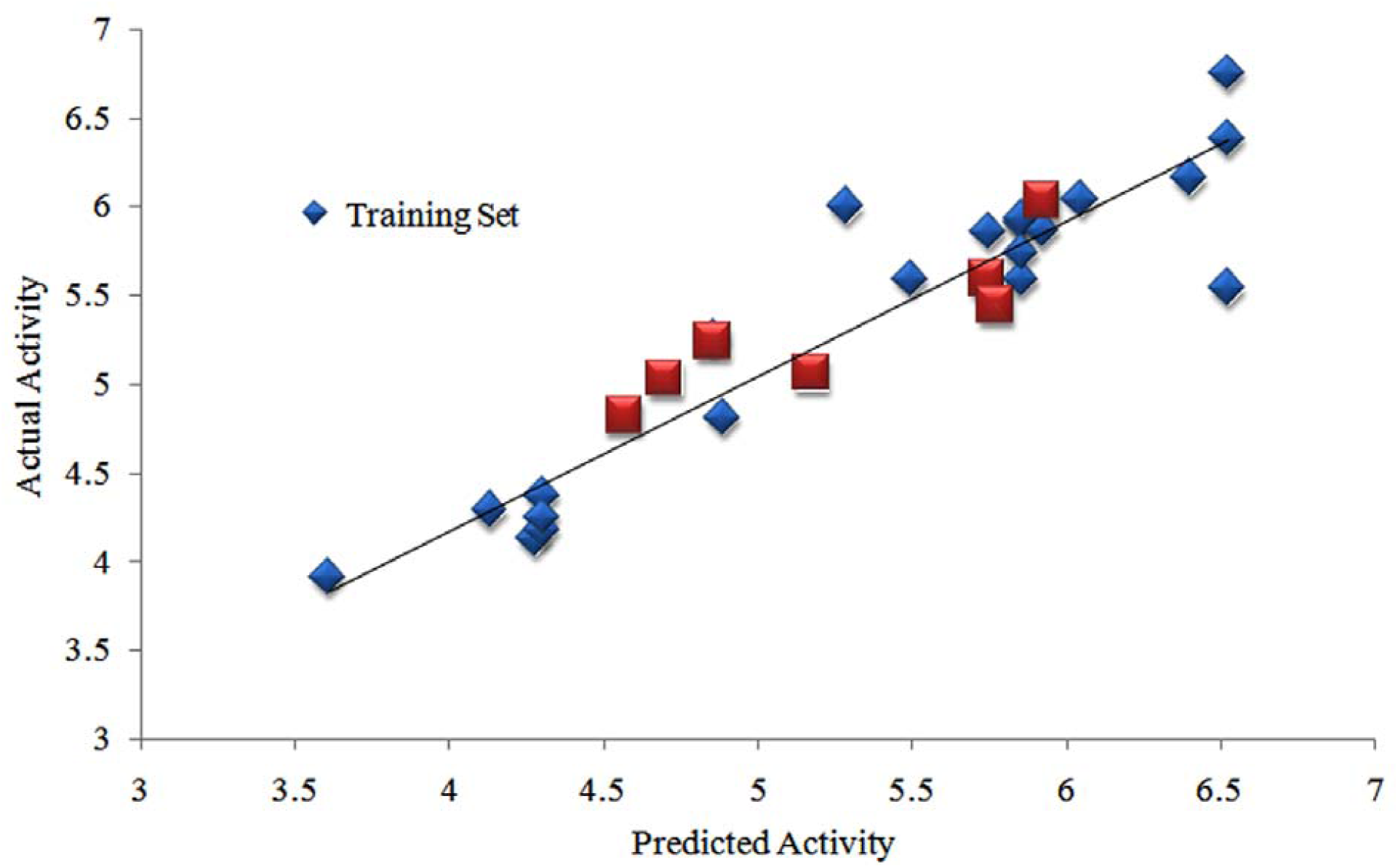
Scatter plot for the predicted and actual pIC50 values for AAAHR hypothesis showing the statistical values of R2 = 0.92, *SD* = 0.16, *F* = 84.8, *N* = 40, *Q*2 = 0.71, RMSE = 0.06, Pearson *R* = 0.90

The statistical parameters R^2^, Q^2^, SD, RMSE and F were used to evaluate the good quality QSAR model. The two models show good and consistent R^2^ greater than 0.82, SD values lesser than 0.4 and F-test values were moderate. This shows that these two models interpreting structure activity relationship for this series of training set compounds satisfactorily. According to Tropsha, high R^2^ is a necessary but not sufficient condition for a QSAR model. Besides the consideration of high R^2^, the best QSAR model should be chosen based on its predictive ability. The AAAHR shows good external predictive ability with Q^2^ value 0.74 and this hypothesis has highest Q^2^ value when compared with other pharmacophore hypotheses, which suggests that AAAHR hypothesis is the best model among the other generated models^**32**^. Additionally, the AAAHR pharmacophore hypothesis has a low RMSE value of the pharmacophore hypotheses, which also supports this hypothesis. Finally, AAAHR hypothesis were considered as the best model based on their R^2^, Q^2^, SD, and RMSE values. The Common pharmacophore modelalignment of active compounds was shown in figure S2. The plots of actual versus predictedpIC_50_in X axis and Y axisin figure 1 for the training set and test set compounds. However, angles and distances of pharmacophore hypothesis AAAHR are given in supplementary material Table S1.

### External Validations of AAAHR pharmacophore model

Established 3D-QSAR AAAHR pharmacophore model were evaluated by external validations with 8 tests set compounds and internally using the leave-n-out technique. The established AAAHR model validated with 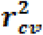 cross validated coefficient produced with the value of 0.56, the high k slope values of regression lines shows the value of 1.002. Non-cross validated correlation coefficient (r^2^) value of 0.51. AAAHR pharmacophore model, possibly will predict new derivatives. The summary of external statistical obtained parameter of 3D-QSAR for the AAAHR pharmacophore model was shown in Table S2.

### QSAR visualization

One of the major advantages of the PHASE 3D-QSAR technique is to get contour cubes based on favorable and unfavorable regions, which could be visualized in 3D space. The contour cubes obtained from AAAHR shows, how 3D-QSAR methods can identify features which is important for the interaction between ligands and their target protein. Such contour cubes allow identification of those positions that require a particular physicochemical property to enhance the bioactivity of a ligand. A pictorial representation of the generated contours was shown in figure 2.In these representations, blue cubes are indicating the sterically favored spatial regions to enhance the activity, whereas red cubes represent the sterically unfavored regions. These cubes can be generated for different properties such as hydrophobic, hydrogen bond donor, hydrogen bond acceptor (electron withdrawing), positive and negative ionic features, which defines the non-covalent interactions with receptor. Visualization of QSAR model cubes associated with hydrophobic property gives an idea about the topology of the receptor site. Figure 2a and 2b shows cubes generated from hydrophobic property using QSAR model while; these cubes illustrate the significant favorable regions and unfavorable hydrophobic/non-polar interactions that arise when the QSAR model is applied to the most active pyridazinone derivative (2-9) and inactive pyridazinone derivatives (2-36).Blue cubes were seen on the most active compound pyridazinone-2-9 at the aromatic ring position. Although comparison of the most significant favorable and unfavorable hydrogen bond acceptors or electron withdrawing features which arise during QSAR model is applied to the most active pyridazinone-2-9 and to the most inactive pyridazinone-2-36 compound was also shown in figure 2a and 2b. Electron withdrawing group (N) and benzene ring structure were associated with blue-colored cubes in the most active compound of pyridazinone-2-9 are observed near atoms which shows high electron withdrawing feature, whereas, in the case of most inactive compound pyridazinone-2-36, red-colored cubes were visible indicating activity difference in these two molecules.

**Figure 2:**
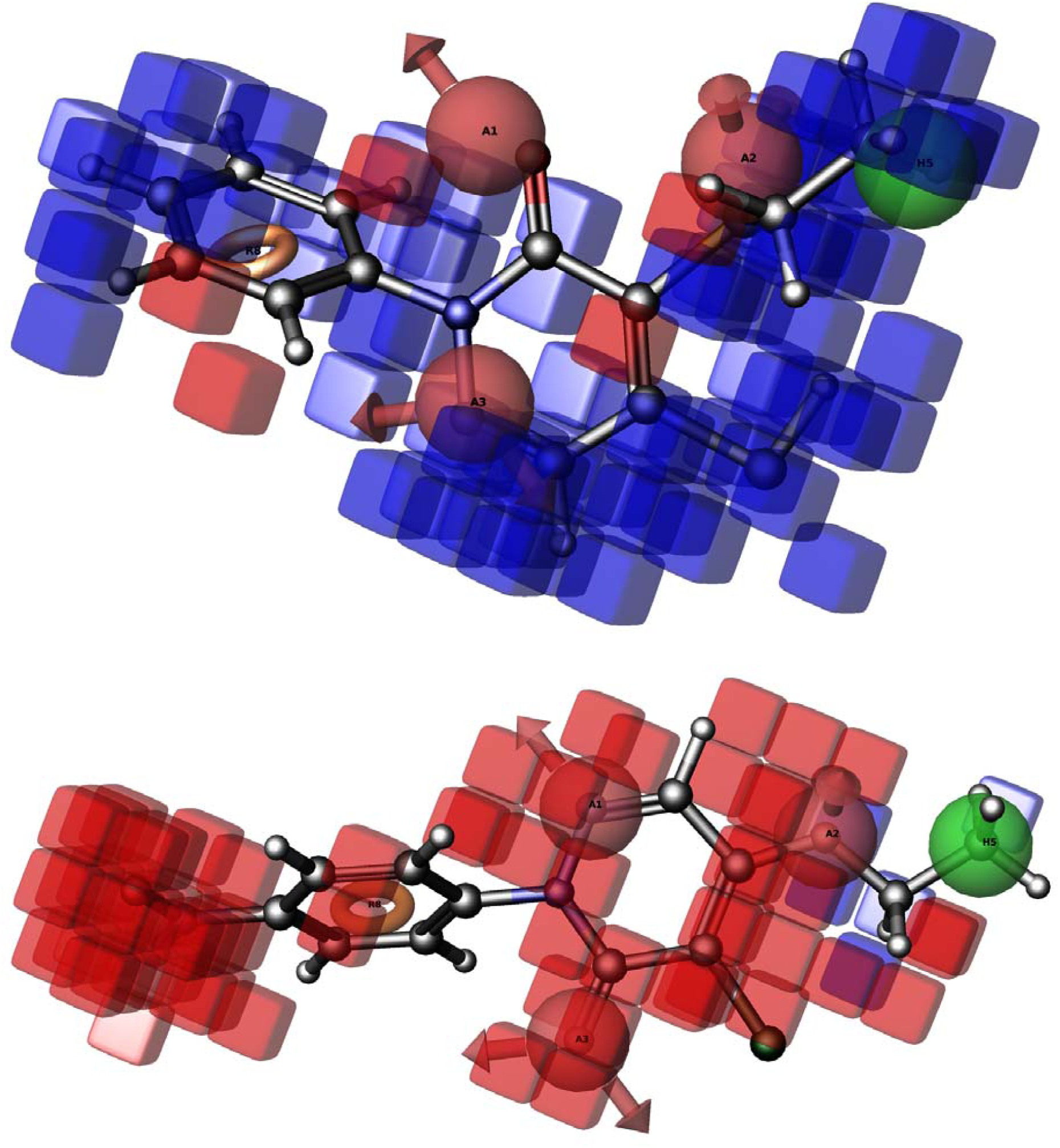
The significant favorable and unfavorable hydrophobic interactions are shown, when QSAR model is applied to the most active compound **(a)** and the most inactive compound **(b)**.

### Validation of Applicability Domain

The validation of AD results represents that the prediction of test set compounds areaccurate and reliable. The DmodX values of eight test set compounds are placed below the critical value of 3.4. The residual values of the predicted vs. experimental SD of X-residuals (DmodX) were represented in figure S3, and it is concluded that these compounds are better for the test set of compounds from this QSAR model. Visualization of the 3D-QSAR model, provides the details of the relationship between structure and activity among these molecules, and thus provides explicit indications for designing the best analogues. Analysis of 3D contour maps was shown to understand the essential structural requirement of the compounds to exhibit better inhibition, which is clearly explained in most active and inactive compounds. While, the successful statistical data based ligand conformations are carried for further study.

### Open and closed lid in the active site of SrtA

The SrtA structure stability, flexibility and intermolecular conformationswere analyzed with this 100ns large scale molecular dynamics simulation. Our previous study on this enzyme for 20ns of simulations has reported about the stability of the protein and so here we tried to extend the simulations for 100ns of time scale. The simulation event involved in this study is to understand the biological happening of the SrtA structure and alterations occurs in each interval of timescale. In the system, the temperature, total energy, mass density, and volume are relatively stable throughout the simulation period, which suggests that the simulations were carried out satisfyingly and system has been well equilibrated. In the figure 3A the RMSD from the NMR structure of C^α^ atoms vs. timescale (ns) shows that initially the protein is more flexible up to 40ns and so the deviations are inconsistent, especially from 2ns to 6ns. The proteins have more deviations due to the presence of loop structures in the protein structure. But after the 40^th^ ns the protein structure was consistent enough as shown in the figure 3. The time evolution of the RMS fluctuation (RMSF) from the averaged structure provides another approach to evaluate the convergence of the dynamical properties of the system as shown in figure 3B.Here, the fluctuations of each amino acid are clearly shown and it’s because of the highly fluctuated loop structure. While RMSF profiles indicate that the residues with higher fluctuation values are those in the long loop Gly53-Asp64 residues (Gly53, Ser54, His55, Met56, Asp57, Ala58, Ser59, Lys60, Ile61, Asp62, Gln63 and Pro64). This particular loop structure was already reported that shown much deviation in *Bacillus subtilis*, and holds the interactions with the lead compounds. In similar to*B. subtilis*, the *Enterococcusfaecalis, Bacillus cereus, Bacillus anthracis, Staphylococcus aureus* are the additional microbial SrtA, which shows the importance of this loop towards the drug-protein interactions^**33-35**^.There was much conformational changes of native protein structure occur in between the 0-100ns. The MD simulation of the SrtA structure showed that the tail like loop structure in the N - terminal region (53-64) present nearby active site which is widely opened at initial position. But this particular loop undergoes more conformational changes at each nanosecond. Therefore, we carefully investigated the structural changes of SrtA with the time interval of every 5ns until 100ns and noticed that the N-terminal tail loop has the biological function of the lid of the active site. The tail loop continuously fluctuating at the dynamic state till the 30ns and after that it slowly changes its position and act as a lid. The lid closes the active site region and it was remains closed till the 100ns. Meanwhile, the conformational changes occur in the SrtA structure was demonstrate in Figure S4 and 4 shows the overall morphed conformations and fig 4 showing the separate conformations of SrtA at each 5ns of interval. The trajectory analysis suggest that loop structure in N-terminal is continuously fluctuating till the 40ns and shows widely opened active site and after 40ns the protein attain its equilibrium condition and loop moderately closes the active site and function as a lid for SrtA structure. Whereas, figure 5 indicates the opening and closing of lid in different timescale of dynamic environment and this lid is called as universal feature of SrtA enzyme in gram positive pathogens. At 40ns the lid closes the active site and function as protector of the active site hydrophobic environment and so it’s difficult task to design the suitable compounds for closed binding pocket of SrtA.The overall simulation studies show good contacts with the system and protein throughout the timescales. In the hydrogen bond analysis between the SrtA and water molecules shows that, there is vast difference in hydrogen bonds. Figure 6 shows the hydrogen bond interactions analysis of SrtA structure with water contact shows that pair above 0.35 nm (A) and pair within 0.35nm (B). The average of hydrogen bond is calculated in two intervals of timescale, i.e. 0-40ns and 40-100ns, as after the 40ns of timescale the loop “lid” has been closed, so it is necessary to understand the interaction difference happen due to this biological function. The average of above 0.35 nm and within 0.35nm wascomparatively high for 0-40ns, compared to after 40ns (40-100ns). The number of hydrogen bonds between the protein and the surrounding water molecules are more in beginning of the timescale (open lid) and comparatively less in later stage of the time scale (closed lid). This is due to closing of active site which protects the hydrophobicity regions in active site to interact with water.

**Figure 3.**
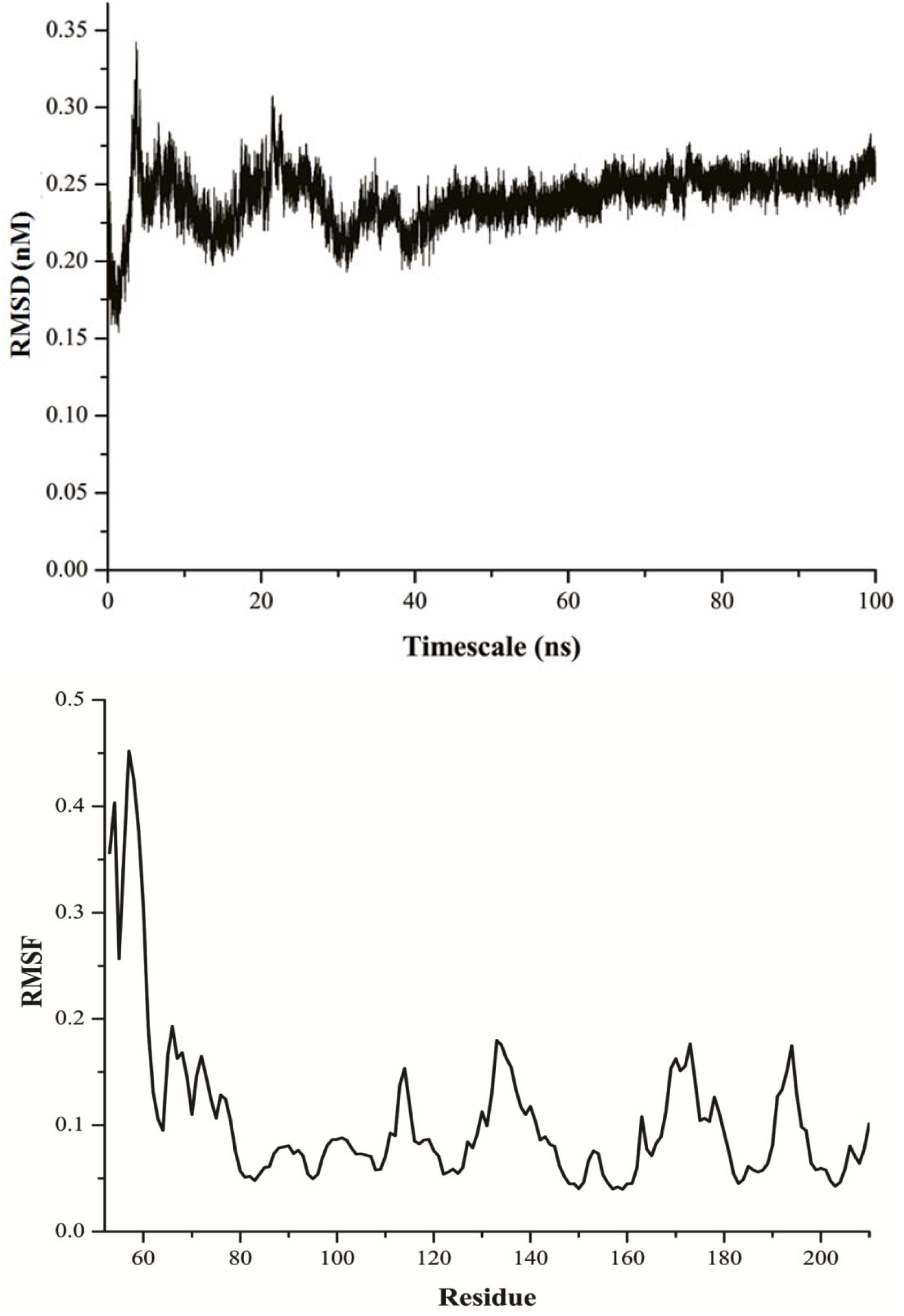
Analysis of SrtA structure in dynamic state with the time scale of 100ns (A) RMSD plot of SrtA for 100ns of timescale. (B) RMSF plot showing residue wise changes in the SrtA structure.

**Figure 4:**
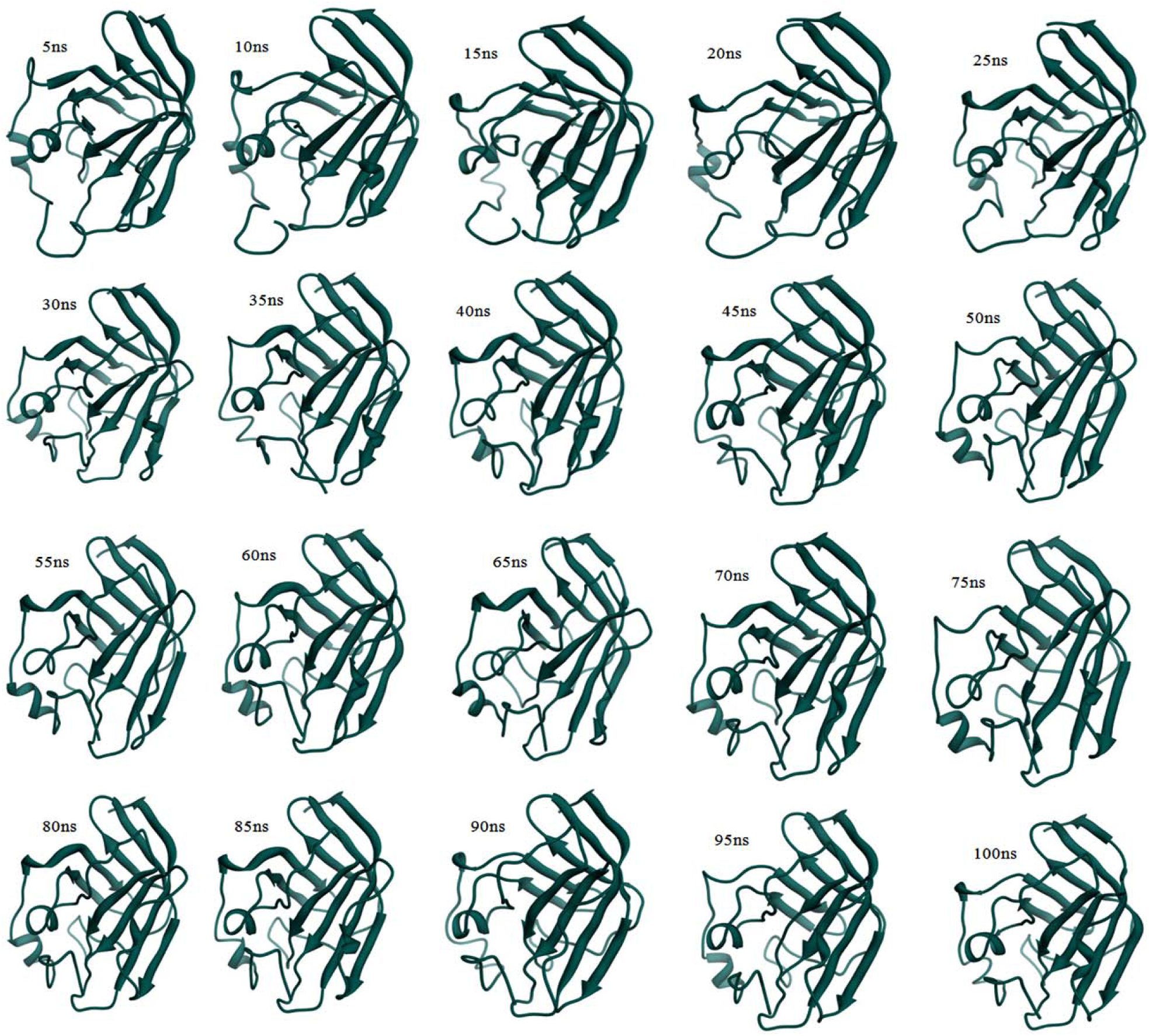
Observation of SrtA active site loop close and open at different timescale (Observed with 5ns interval)

**Figure 5:**
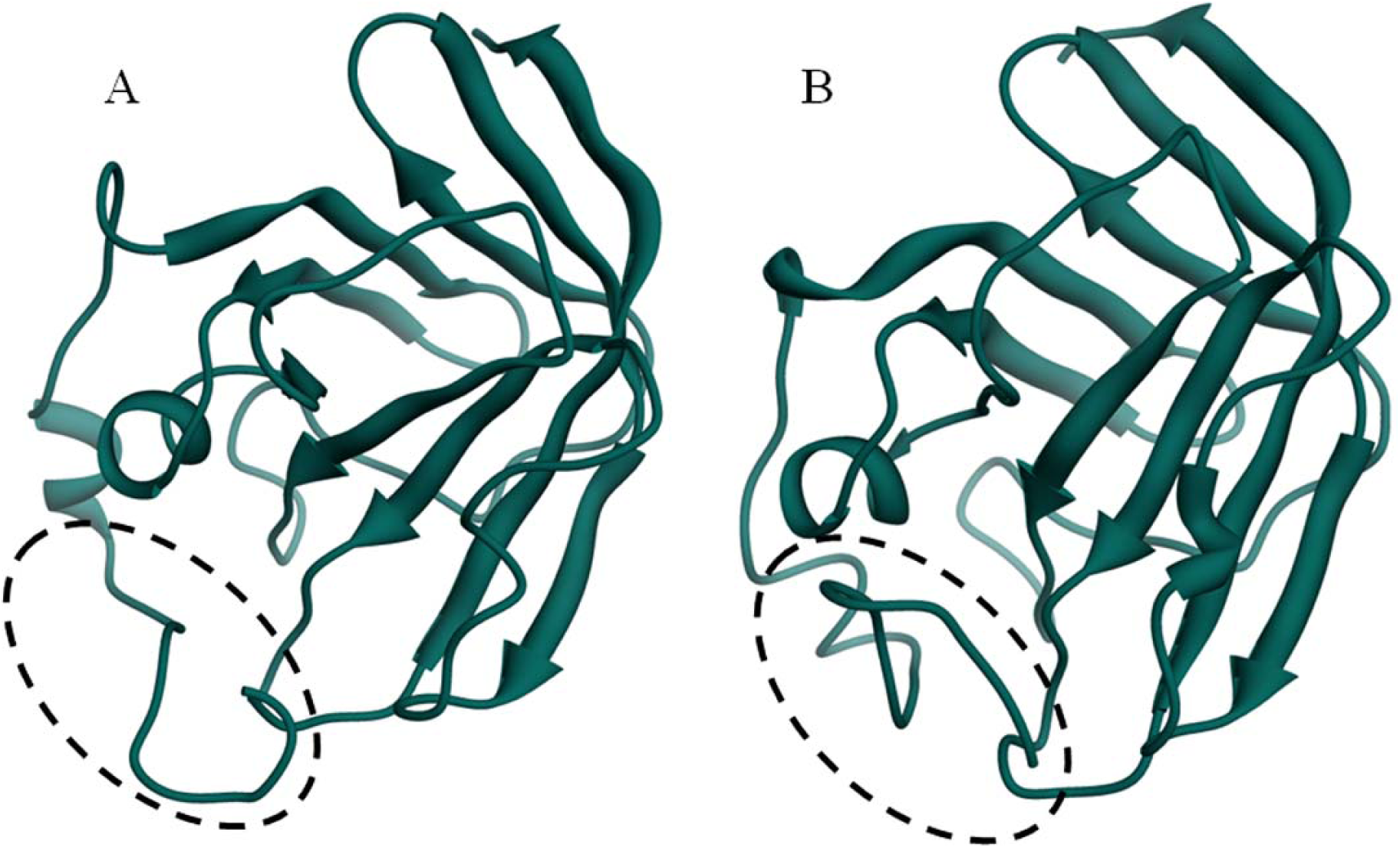
Active site loops on dynamic state (A) showing the active site lid in open state at 0ns (B) Active site lid closed at 40ns of timescale

**Figure 6.**
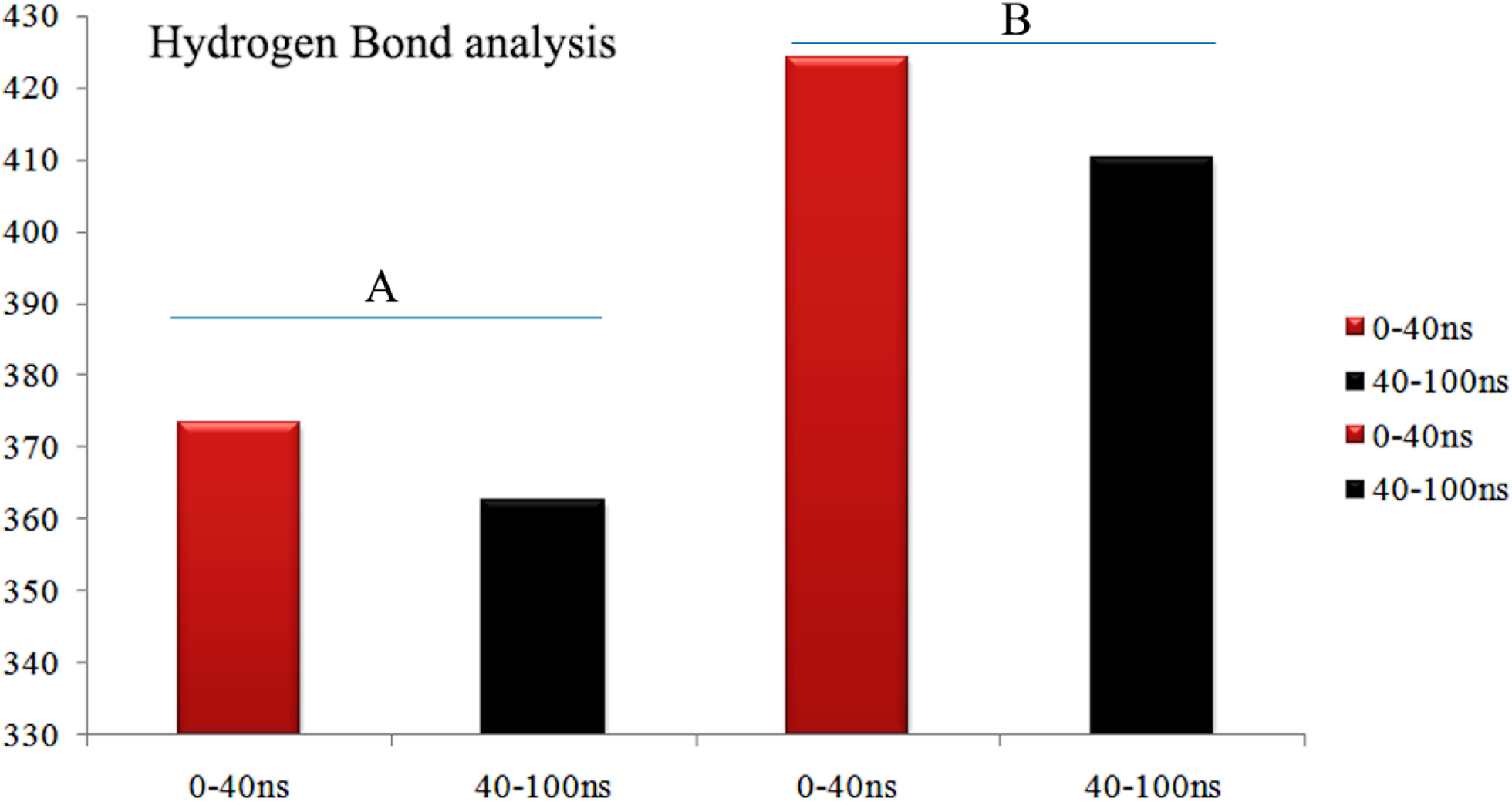
Hydrogen bond analysis of SrtA structure with water contact shows that pair above 0.35 nm (A) –and pair within 0.35nm (B) - are showing high contacts in 0-40ns and after that the hydrogen bonds has been decreased.

### SrtA inhibitors against Open and Closed lid conformations

This research works, explains the statistical validation and confirmation of SrtA inhibitors using closed and open lid of SrtA structure through molecular modeling techniques. In *Staphylococcus aureus*, the substrate amino acid Leu plays the crucial role of an anchor to the loop between the beta 6-beta7. This interaction directing the conformational transition of the enzyme from an “open” to a “closed” state subsequent to which the Pro residue facilitates the consummation of binding through predominant engagement of the loop and catalytic motif residues in hydrophobic interactions. Through this, the SrtA loop function in *B. anthracis* has been clearly explored with the closedand open active site conformations taken from the molecular dynamics simulations. In order to check active conformation of SrtA the bioactive conformers of the inhibitors obtained from the 3D QSAR are validated in both open and closed lid conformations. The QPLD and LSBD (ligand and structure based descriptors) calculations of both closed and open lid conformations were used for obtaining the values of activity based on the conformations of the proteins. The docking procedures used this study aim to identify correct poses of ligands in the binding pocket of a protein and to predict the affinity between the ligand and the protein. LSBD calculations predict the activity based on docking pose with respect the experimental activity. The purpose of the scoring procedure is the identification of the correct binding pose with lowest energy value, and the ranking of protein-ligand complexes based on their binding affinities. So that, bioactive conformations obtained from 3D QSAR are made rigid conformations and both closed and open lid are interacted. The analysis of binding patterns from the docking suggests that closed binding pocket has three main amino acids (Met56, Val110 and Trp171), which are mainly involved in hydrophobic interactions. Almost 92% of the active compounds show the interactions with this catalytic triad of SrtAand compound pyridazinone-2-17 is additionally showing the hydrophobic interactions with Ala58. As discussed earlier, the lid closed the binding pocket which protects the hydrophobic surface of the active site and these experimentally validated compounds are well suited for the closed lid binding pocket of SrtA. From the interactions, it is concluded that the aromatic group present in the pyridazinone and pyrazolethione are core important for the activity. The figure 7 shows the ligand aromatic ring structure involved in the hydrophobic interactions with Met56, Val110 and Trp171. The hydrophobic interactions represented in green colored dashed lines in Figure 7 indicate the hold of aromatic ring structure inside the hydrophobic binding pocket. Among the interacting amino acids Met56 is contributing in the function of the closed lid loop and also it’s actively participated in the hydrophobic interactions. This result indicates that those three amino acids are core important for targeting SrtA enzyme and the aromatic ring structure of pyridazinone and pyrazolethione are play a key role in inhibition of SrtA activity. Best conformations of SrtA inhibitors, shown in figure 7, with both protein conformations are taken into account for the Molecular Dynamics Simulation.

**Figure 7:**
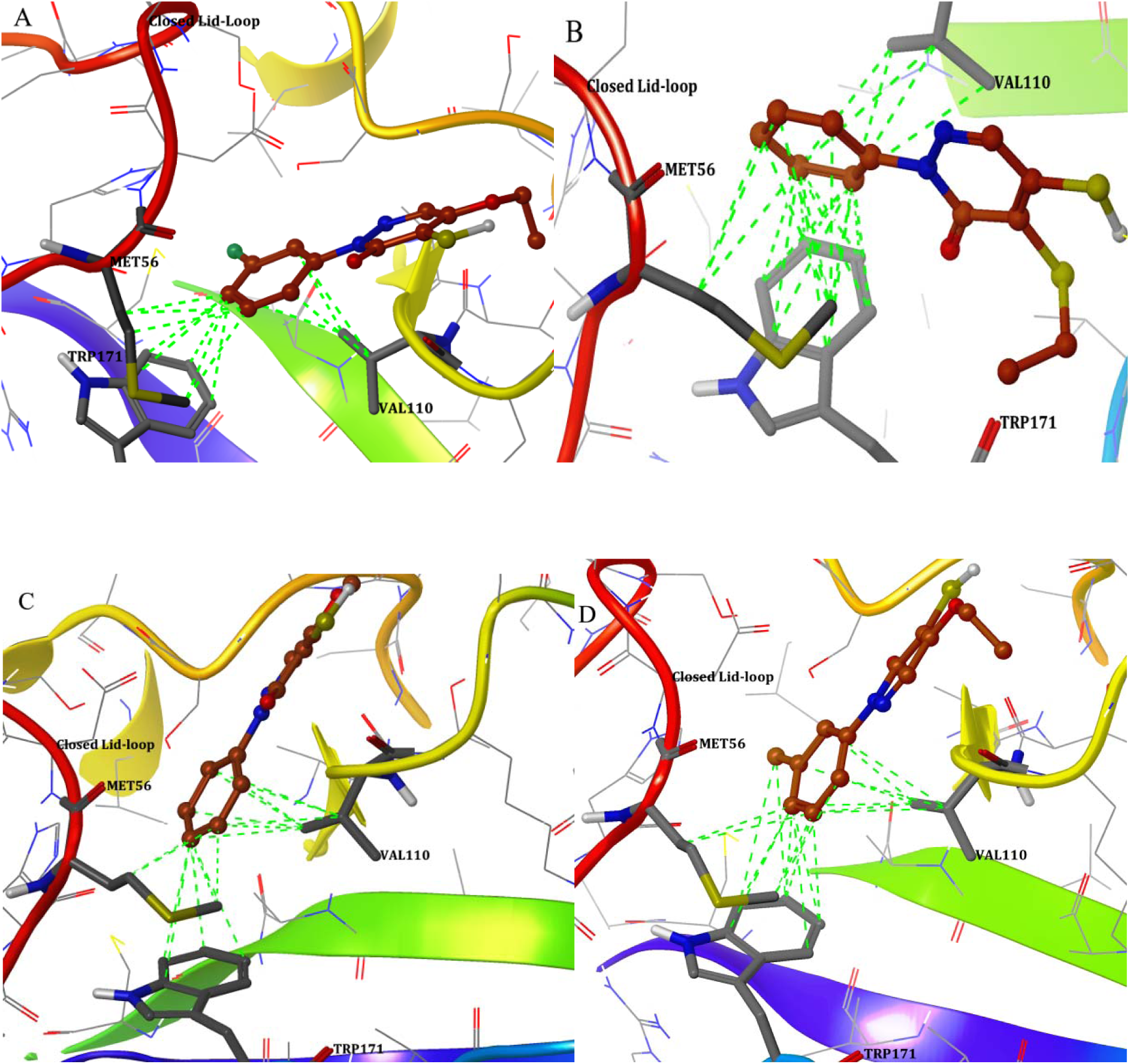
Active compounds showing the aromatic ring structure showing hydrophobic interactions with Met56, Val110 and Trp171

### Ligand Stability-Molecular Dynamics Simulation

Based on activity, docking and binding energy, five best ligands were chosen for molecular dynamics simulation studies. In the analysis of trajectories shows that all the compounds on this dynamic event having better interaction throughout the timescale of 20ns and ligand does not detach from the protein. The figure S5 shows the RMSD graph of top five active compounds and these compounds are more stable inside the binding pocket due to the engulf of the hydrophobic interaction with the aromatic ring. Moreover, the figure explained that experimentally validated compounds have stable interaction and bind properly inside the closed binding pocket of SrtA. In the initial state all the compounds is slightly moved from its original position and after that the ligand binding position got stable inside the binding pocket and the hydrophobic bonds holds the ligand strongly by not allowing the ligands to detach from the protein structure. Depth analysis of trajectories clearly shows that the catalytic triad of Met56, Val110 and Trp171 are playing role in holding ligand and also interacted with aromatic ring. The average mean values of RMSD for all test compounds (pyridazinone-2-19, pyridazinone-2-9, pyrazolethione-3-12, pyridazinone-2-5 and pyridazinone-2-20) were calculated and those values clearly shows that all are having the average values of less than 0.2nm to 0.4nm. Pyridazinone-2-5 is having 0.4nm average RMSD, which is comparatively higher than the other four compounds. When analyzing RMSD values and graph, all the experimentally verified compounds are stable and have much correlation with each other. Interestingly, all the compounds are having the same binding modes in dynamic state and provides more similar pattern of inhibition. These results suggested that binding of the ligand to the protein showed a deviation from their initial position because of adjustments in their configuration, but remain bound within the catalytic triad of the protein.

### Energy Calculation Accounts for Activity Prediction

In general most of the studies measure the accuracy of scoring function by their ability to correctly rank the activity of a congeneric set of ligands. The ligand activity prediction against both open and closed lid conformation of SrtA is equally important in drug design of *B. anthracis*. The experimental activity reported by Suree*et al*., 2009 is considered in pIC_50_ and liaison calculation is analyzed, while the liaison is a method of predicting ligand-protein binding free energies using a model that has been correlated to known binding energy values. The process involves two steps, a fitting step and a predicting step and each step are carried out as a two tasks, which are the simulation task and an analysis task. The experimental binding affinities given in the property activity (kcal/mol) will be used for the response or dependent variable from which a linear model will be created. The LIA equation outlines in detail and it estimate binding affinities using <U_ele_>, <U_vdw_> and <U_cav_> terms from Liaison. Once the LIA model has been created, it must be analyzed to see if it makes intuitive sense and possesses the predictive power that does not arise by chance. Here, the obtained energy levels are analyzed with their reference activity and theoretical activity of both closed lid and open lid ligand binding complex. The obtained activities of experimental and theoretical values are plotted and R^2^ is calculated to understand the correlation between the both with respect to open and closed lid. The correlation analysis shows in the figure 8, predicts that, the open lid protein conformations shows *r* ^2^ of 0.730 and closed lid conformations shows *r*^2^ of 0.925. This confirms that, the protein conformation obtained from the 40^th^ ns of molecular dynamics simulation is more active and suitable for molecular modeling calculations.

**Figure 8.**
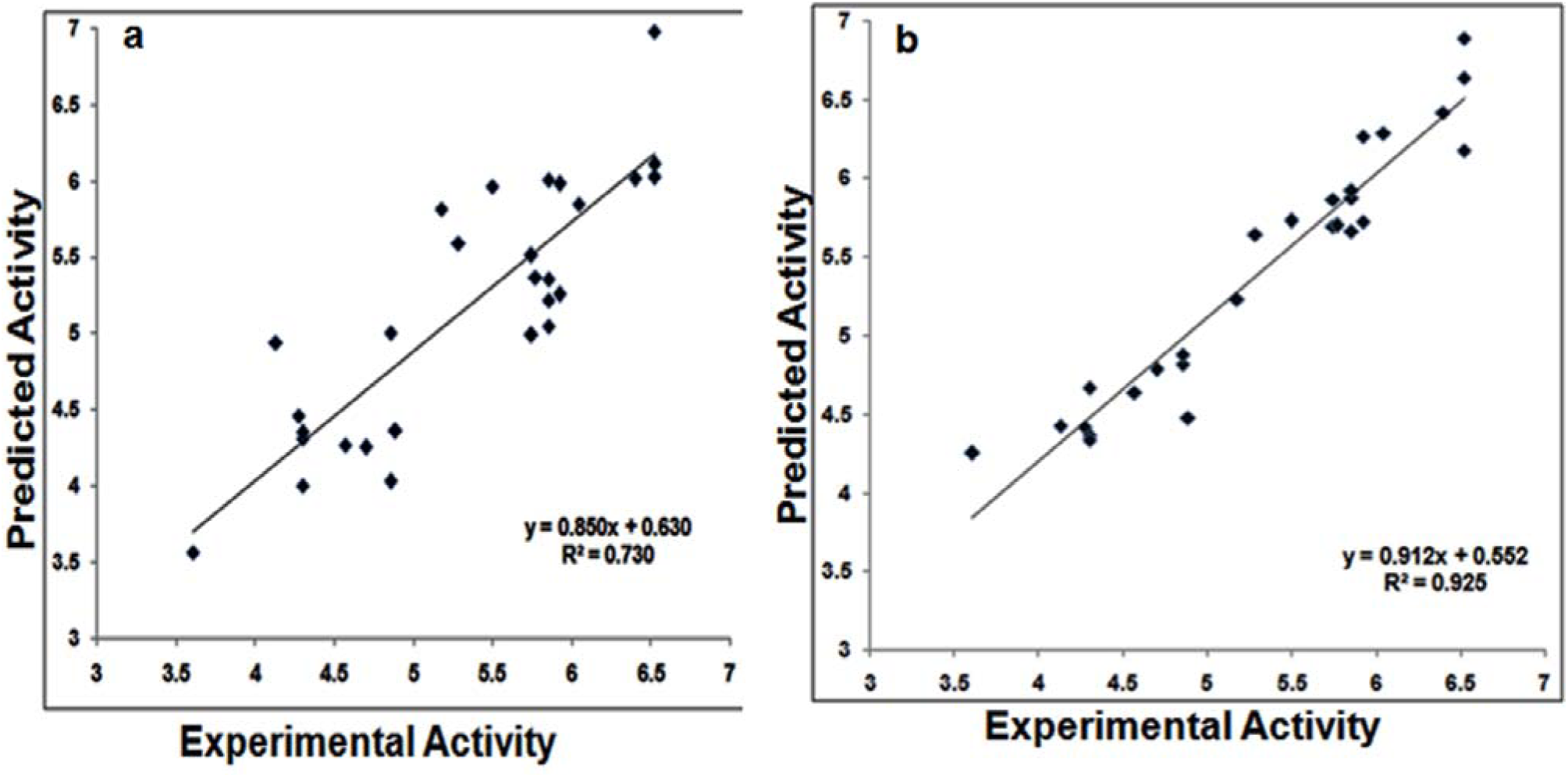
R^2^ cross validation of theoretical and experimental activity with respect to open lid (a) and closed lid (b) conformation

## Conclusion

In this current study, 3D pharmacophoric features and QSAR of the selected SrtA inhibitors were explained, which are experimentally validated against the *B. anthracis*SrtA. The pharmacophore with AAAHR pharmacophoric features areassociated with a 3D-QSAR model giving the good statistical significance and predictive ability. The 3D contour map analysis was helped to understand the essential structural requirement of the compounds to exhibit the better inhibition, which is clearly explained in the most active and inactive compounds. Meanwhile, the N-terminal tail loop is playing role as the protector lid of the active site hydrophobic regions was through molecular dynamic studies. Through MD simulation, we reported that open lid conformations have a mechanism of closed lid active site at 40ns. Furthermore, the activity prediction of the experimentally proven SrtA inhibitors was analyzed by docking the selected SrtA inhibitors into both conformations of open and closed lid of active site through QPLD method. Although the open lid protein conformation shows *r* ^2^ of 0.730 and closed lid conformation shows *r* ^2^ of 0.925. This confirms that, the conformation obtained from the 40^th^ ns of molecular dynamics simulation is more active and suitable for molecular modeling calculation. The interaction between ligand and protein are observed for catalytic triad Met56, Val110 and Trp171 by means of hydrophobic interaction. From these studies, it is understood that the aromatic ring present in the rhodanine, pyridazinone and pyrazolethione derivatives are playing an important role in the inhibition. Consequently, the ligand bound complexes are confirmed and selected compounds have stable interaction by holding the catalytic triad in binding region. Finally, atom-based 3D-QSAR model, molecular dynamics and docking studies performed here could be very valuable for the development of new and potent lead compounds for the inhibition of SrtA.

